# Optimizing the Timing and Composition of Therapeutic Phage Cocktails: A Control-theoretic Approach

**DOI:** 10.1101/845172

**Authors:** Guanlin Li, Chung Yin Leung, Yorai Wardi, Laurent Debarbieux, Joshua S. Weitz

## Abstract

Viruses that infect bacteria, i.e., bacteriophage or ‘phage’, are increasingly considered as treatment options for the control and clearance of bacterial infections, particularly as compassionate use therapy for multi-drug resistant infections. In practice, clinical use of phage often involves the application of multiple therapeutic phage, either together or sequentially. However, the selection and timing of therapeutic phage delivery remains largely ad hoc. In this study, we evaluate principles underlying why careful application of multiple phage (i.e., a ‘cocktail’) might lead to therapeutic success in contrast to the failure of single-strain phage therapy to control an infection. First, we use a nonlinear dynamics model of within-host interactions to show that a combination of fast intra-host phage decay, evolution of phage resistance amongst bacteria, and/or compromised immune response might limit the effectiveness of single-strain phage therapy. To resolve these problems, we combine dynamical modeling of phage, bacteria, and host immune cell populations with control-theoretic principles (via optimal control theory) to devise evolutionarily robust phage cocktails and delivery schedules to control the bacterial populations. Our numerical results suggest that optimal administration of single-strain phage therapy may be sufficient for curative outcomes in immunocompetent patients, but may fail in immunodeficient hosts due to phage resistance. We show that optimized treatment with a two-phage cocktail that includes a counter-resistant phage can restore therapeutic efficacy in immunodeficient hosts.

## 1 Introduction

The spread of multi-drug resistant (MDR) pathogens is a global public health crisis [O’neill, 2014]. The threat of the antibiotic resistance crisis has spurred research and development of alternative antimicrobials, including phage [Kortright et al., 2019, Chan et al., 2013, Kutter et al., 2015, Young and Gill, 2015, Merril et al., 2003, Chan et al., 2016, 2018, Schooley et al., 2017, Forti et al., 2018, Dufour et al., 2019]. Phage has been applied in compassionate use scenarios, for example, to successfully cure patients both in the USA and in Europe [McCallin et al., 2019, Dedrick et al., 2019, Chan et al., 2018, Schooley et al., 2017, Jennes et al., 2017], catalyzing the 2018 launch of the first North American phage therapy center based at UCSD (IPATH). Yet, despite individual successes, phage therapy has a mixed record in controlled clinical trials. For instance, a large-scale trial involving more than 200 patients failed to demonstrate that phage treatment improved outcomes for children infected by Escherichia coli with symptoms of severe diarrhea in Bangladesh [Sarker et al., 2016]. Similarly, the recent European phase II clinical trial to treat burn wound patients failed to show superiority compared to a reference treatment [Jault et al., 2019].

The host immune response is an important driver of within-host infection dynamics, but the aforementioned phage therapy studies have not considered the effects of host immune status in shaping the outcomes of phage therapy. To address this question, we have developed mathematical models in prior work that consider the tripartite interactions between phage, bacterial pathogen, and host innate immunity [Leung and Weitz, 2017]. Combined with animal experiments, the results have shown that bacterial populations are not necessarily eliminated by either phage or the immune response alone. Instead, bacteria are eliminated when phage and the immune response work in synergy [Roach et al., 2017]. Importantly, curative success was not inevitable. For example, phage therapy was ineffective in innate immune activation deficient hosts and neutropenic hosts. Therapeutic failure was caused by the spread of phage-resistant bacteria as predicted by the mathematical models. Such failure raises a new challenge: is it possible to rationally combine phage strains, dosage, and targeting to overcome therapeutic failure in immunodeficient hosts and in other scenarios such as rapid phage clearance from the host?

Control theory is a potentially useful approach to address the problem of optimizing the dosage, timing, and composition of therapeutic agents. For example, control theory has been applied to optimize antiretroviral drug therapy for HIV infections [Culshaw et al., 2004, Jang et al., 2011, Croicu, 2015, Croicu et al., 2017], minimize resistance in antibiotic treatment [Peña-Miller et al., 2012], and determine the optimal dosing schedule of antimalarial medications [Thibodeaux and Schlittenhardt, 2011] and cancer therapies [Castiglione and Piccoli, 2006, de Pillis et al., 2008, Ledzewicz et al., 2012]. These applications of control theory have focused on modeling the within-host disease dynamics using a set of coupled nonlinear differential equations describing the population dynamics of disease agents such as pathogens or tumor cells, as well as host cells that include immune cells and/or cells targeted by pathogens. The cost function to be minimized is then chosen to balance a number of treatment goals, including minimizing pathogen/tumor load, maximizing healthy cell populations, and limiting treatment costs and toxicity.

As examples beyond within-host treatments of diseases, control theory has also been applied to optimize strategies in controlling between-host transmission of infections. In these studies, the spread of the infectious disease is modeled by standard epidemiological models such as the Susceptible-Infected-Susceptible (SIS) model or the Susceptible-Infected-Recovered (SIR) model. The control strategies consist of epidemiological interventions such as vaccination, sanitation, and treatment of infected individuals. The cost function is determined based on minimization of the infected population or number of deaths subjected to costs of the control efforts. Such epidemiological applications of control theory have been used to optimize control strategies in vector-borne diseases [Blayneh et al., 2009], cholera epidemics [Neilan et al., 2010], anthrax infection in animals [Croicu, 2019], and infectious disease with two strains of pathogens [Rowthorn and Walther, 2017].

In this paper, we develop a control-theoretic framework to optimize monophage therapy and multiphage (cocktail) treatment of immunodeficient hosts and in other scenarios where standard phage therapy is likely to fail. In Sect. 2, we introduce the mathematical model of phage therapy and define the control problem. In Sect. 3, we analyze the optimal control problem, show that the optimal control solution exists, and derive the necessary conditions for the optimal control via Prontryagin’s maximum principle [Pontryagin et al., 1962]. Then, we implement a Hamiltonian-based algorithm [Hale et al., 2016, Wardi et al., 2016] to numerically compute the optimal control solutions for monophage therapy and a phage cocktail treatment consisting of two phage strains. The numerical results are presented in Sect. 4 followed by a discussion in Sect. 5.

## 2 Problem Formulation

### 2.1 A Mathematical Model of Phage Therapy

We propose a phage therapy model that considers the nonlinear dynamics arising from interactions between sensitive bacteria *S*, phage-resistant bacteria *R*, phage *P_S_* (i.e., only targeting sensitive bacteria), phage *P_R_* (i.e., only targeting phage-resistant bacteria) and the host innate immune response *I*, see Fig. 1. Hence, this model intentionally makes the assumption that phage are specialized in their infection of bacteria; generalizations of this approach are considered in the Discussion. Two strains of bacteria (*S* and *R*) reproduce given the limited environmental capacity *K_C_*. The phage-resistant bacteria are emerged from sensitive bacteria through mutation with a fixed probability *μ* per cellular division. Both strains of bacteria are killed by the immune response and both stimulate immune activation. The immune response is stimulated by the presence of bacteria with a maximum activation rate *α* until it reaches the maximum capacity *K_I_*. Phage populations *P_S_* and *P_R_* infect and lyse sensitive and phage-resistant bacterial populations at rates *F*(*P_S_*) and *F*(*P_R_*) respectively. In this study, we assume that the two phage types (*P_S_* and *P_R_*) have identical adsorption rate *ϕ*, burst size *β* and decay rate *ω* for simplicity. In general, these two trait parameters can be strain-specific, and the optimal control analysis and simulation procedures will be the same. The treatments inject phage into the system, the injection rates of phage *P_S_* and phage *P_R_* at time t are *ρ_S_* (*t*) and *ρ_R_*(*t*) respectively. The dynamics of bacteria, phage and the innate immune system can be modeled using the following system of nonlinear differential equations,

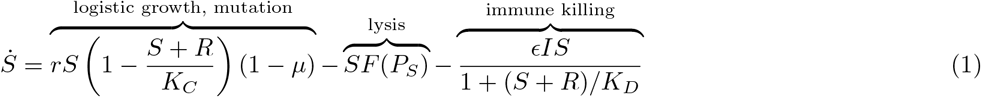

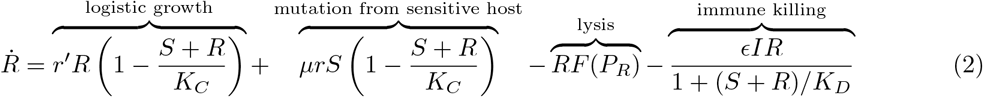

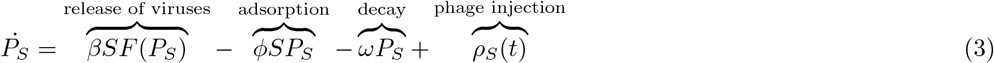

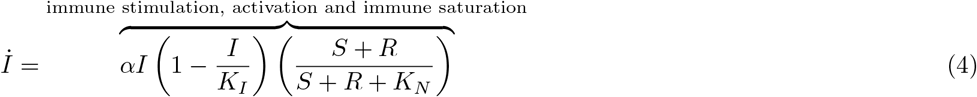

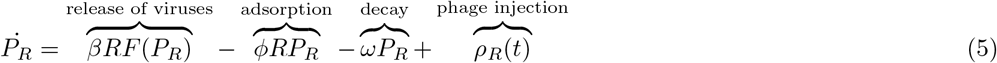

where 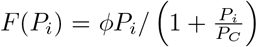 for *i* ∈ {*S,R*} is the phage infection rate that characterizes the effect of phage saturation during the infection. Specifically, phage saturation occurs when multiple phage adsorb to the same target bacterial cell when at high phage population density. We constrain the injection rates of two phage doses as following,

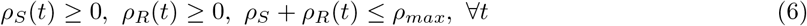

where *ρ_max_* is the fixed maximal injection rate. The goal is to control the phage injection rates *ρ_S_*(*t*) and *ρ_R_*(*t*) so as to minimize the bacterial population over the entire treatment and at the final time, while limiting the amount of phage injected into the body (*i.e*., treatment costs). The specific cost functional associated with this goal will be introduced later in Sect. 2.2.

**Fig. 1.**
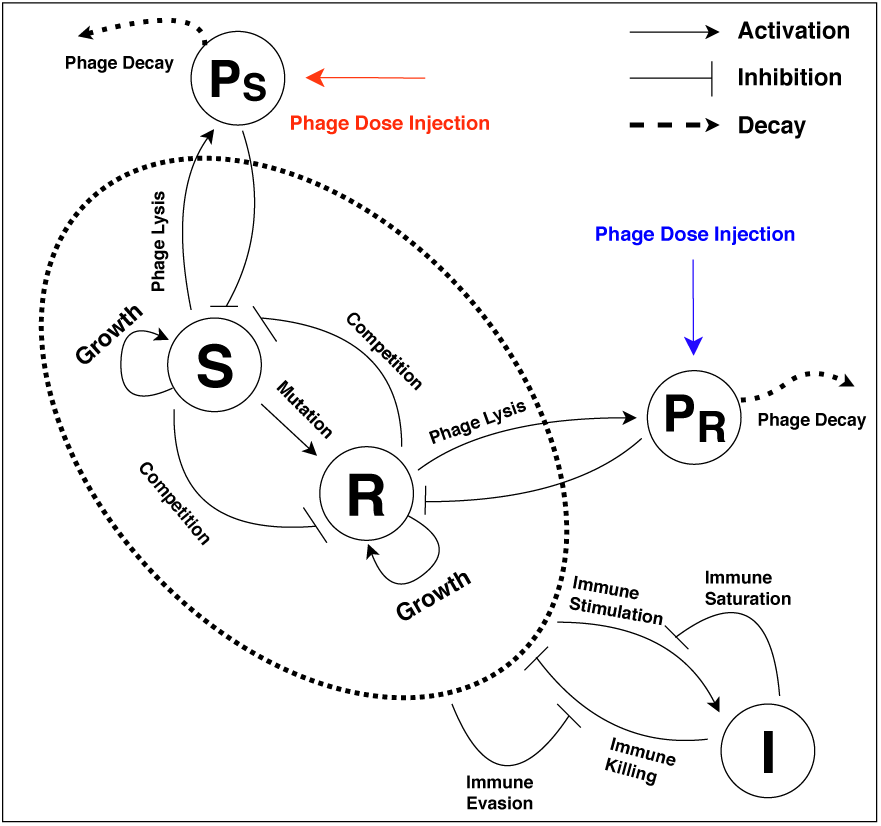
Schematic of phage therapy model in the system (1)–(5). Sensitive bacteria (*S*) and phage-resistant bacteria (*R*) are targeted by phage (*P_S_*) and phage (*P_R_*), respectively. Innate immunity (*I*) is activated by the presence of bacteria and attacks both bacterial strains.

#### Scaled Model

To simplify the system (1)–(5), we rescale state variables and parameters. As the state variables and parameters have different units, we transform them, by scaling, into non-dimensional variables. Accordingly, the dimensionless state vector *x* is

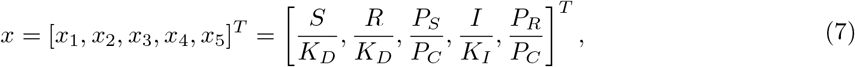

where (·)^*T*^ is the notation of matrix transpose. The scaled model parameters and control variables are:

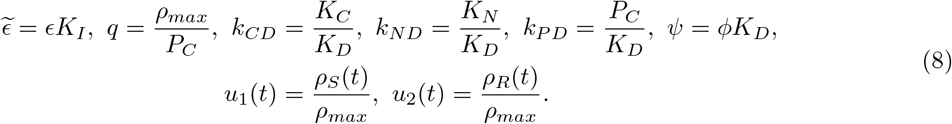

In doing so, the resulting scaled system is

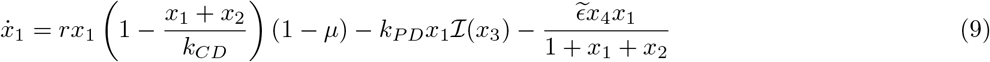

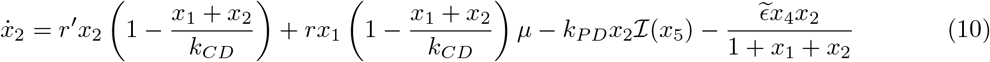

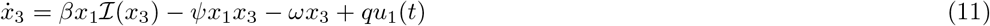

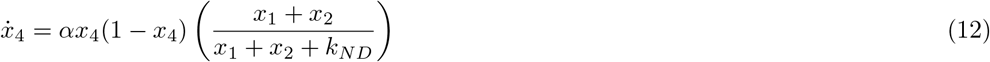

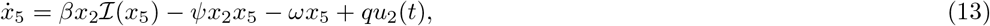

where 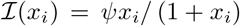 for *i* ∈ {3,5} is the scaled phage infection rate function. Note that the injection rates (*ρ_S_,ρ_R_*) are scaled by the maximal injection rate *ρ_max_*, the scaled injection rates (*u*_1_, *u*_2_) are interpreted as the relative intensity (or strength) of the maximal injections, we have constrained: *u*_1_(*t*) ≥ 0, *u*_2_(*t*) ≥ 0 and *u*_1_(*t*) + *u*_2_(*t*) ≤ 1, ∀*t*.

### 2.2 Objective Functional

Define the set *U* ⊂ ℝ^2^ as a convex and compact set, 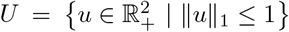 where 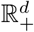 is the *d* dimensional non-negative orthant. We define the space of *admissible controls*, denoted by 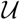, as the set of Lebesgue-measurable functions *u*: [*t*_0_, *t*_*f*_] → *U*. The cost functional to be minimized over 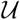 is written in the following Bolza form,

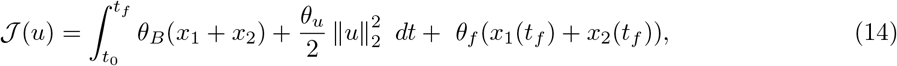

where *θ_B_, θ_u_* and *θ_f_* are the regulator weights. We denote 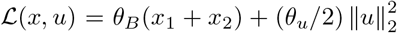 as the *running cost*, which is the integrand of 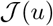. The *terminal cost* is denoted as *g*(*x*(*t_f_*)) = *θ_f_*(*x*_1_ (*t_f_*) + *x*_2_(*t_f_*)). The optimal control problem is

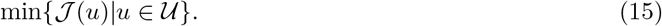

This optimal control problem may not have a solution in the sense that the minimum in Eq. (15) does not exist. But the infimum exists, and we denote it by

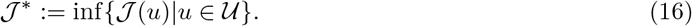

Moreover, suppose the minimizer in Eq. (15) exists (we will prove the existence in Sect. 3.1), it may be impractical to implement it as a control unless *u*(*t*) is piece-wise continuous in *t* (by definition of the problem, it only needs to be Lebesgue measurable). Therefore, we resort to a numerical algorithm designed to compute a piecewise-continuous control *u* such that 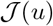 approximates 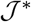 within any pre-set degree of precision, the algorithm description can be found in Sect. 4.1.

## 3 Analysis of Optimal Controls

### 3.1 Preliminaries

#### Positively Invariant Set

For many complex population dynamics models, the populations remain bounded forward in time, e.g., virus-host interactions [Beretta and Kuang, 1998], vector-borne diseases [Blayneh et al., 2009]. The following proposition guarantees the boundedness of system (9)–(13) for any controls 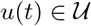.

##### Proposition 1

*Let Ω be the following subset of* 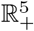:

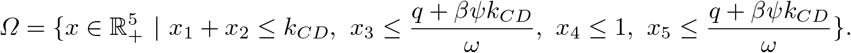

*Then, Ω is a positively invariant set under system (9)–(13)*.

*Proof*. It should be clear that the state solutions are bounded from below by zero such that 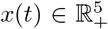 for all *t* ≥ 0, i.e., the densities of bacteria, phage and immune cells cannot be negative. The following discussion assumes the initial condition is in set *Ω, i.e., x*(0) ∈ *Ω*. Note that

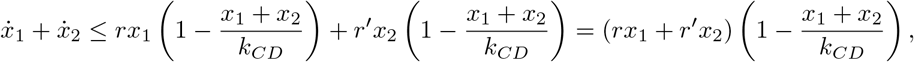

which implies that *x*_1_ (*t*) + *x*_2_ (*t*) ≤ *k_CD_* for *t* ≥ 0. Similarly, we must have *x*_4_(*t*) ≤ 1 for all *t* ≥ 0. The control inputs (*u*_1_, *u*_2_)^*T*^ ∈ *U* for all *t* ≥ 0, hence, we have

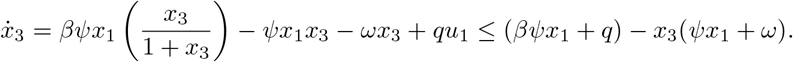

Using the boundedness of *x*_1_, *i.e*., 0 ≤ *x*_1_ ≤ *k_CD_*, we have

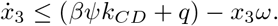

Clearly, *x*_3_(*t*) ≤ (*q* + *βψk_CD_*)/*ω* for all *t* ≥ 0. Similarly, using the boundedness of *x*_2_, we have *x*_5_(*t*) ≤ (*q* + *βψk_CD_*)/*ω* for all *t* ≥ 0. Altogether, we find that *Ω* is positively invariant under system (9)–(13).

#### Existence of Optimal Control

Notably, the system (9)–(13) is control affine and the control set 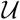 is compact and convex, the integrand of the cost functional, 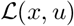, is convex on *U* for each *x*, and the terminal cost function *g*(*x*) is continuous. Thus, the sufficient conditions for the existence of optimal control are satisfied (see Theorem 4.1 in [Fleming and Rishel, 1976]). The similar exercises of proving the existence of optimal control based on Theorem 4.1 in [Fleming and Rishel, 1976] are referred to [de Pillis et al., 2008], [Jang et al., 2011], [Camacho et al., 2014], and [Blayneh et al., 2009].

### 3.2 The Optimality System

The existence of optimal control has been established, we derive the optimality system by Pontryagin’s Maximum Principle (PMP). Note that PMP gives the necessary conditions for the optimal control [Pontryagin et al., 1962]. First, we formulate the optimal control problem as following

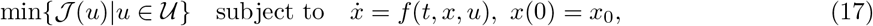

where 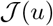 is the cost functional given in Eq. (14), 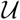 is the admissible control space, *f*(*t,x,u*) is right hand side (RHS) of system (9)–(13) and *x*_0_ is the initial condition. Applying PMP, we obtain the optimality conditions that must be met for an optimal control in problem (17).

#### Theorem 1

*If* 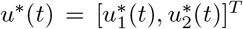 *is an optimal control pair for the control problem in Eq. (17), x*(t) and λ*(t) are the corresponding state trajectory and costate trajectory, then*

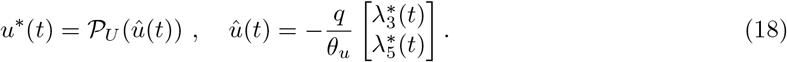

*where 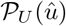 represents the projection of û onto U in ℓ*_2_-*norm. The detailed implementation of projection operator 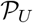 is given in Appendix A*.

*Proof*. Given that 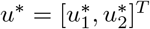 is an optimal control pair for the control problem in Eq. (17), and *x*\*(*t*) and *λ*\*(*t*) are the corresponding state trajectory and costate trajectory, then the following equations are satisfied by PMP:

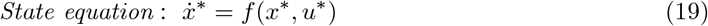

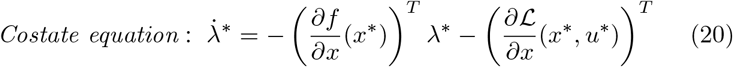

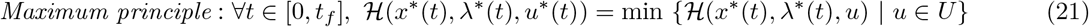

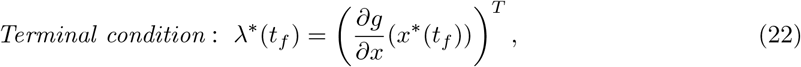

where 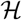 is the Hamiltonian with form of 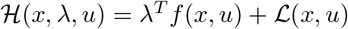. In our case, we have

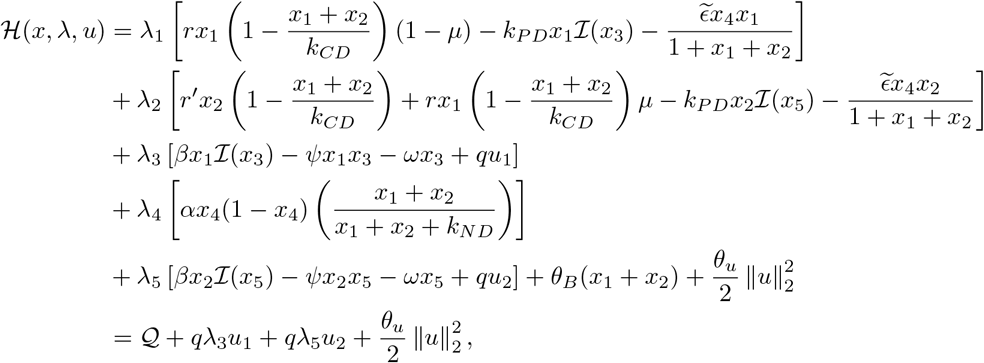

where 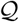 is the collection of terms that has no argument in *u*. The minimization of 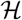 over *u ∈ U* is a linear constrained quadratic programming (QP) problem. We write the minimization problem in its equivalent form,

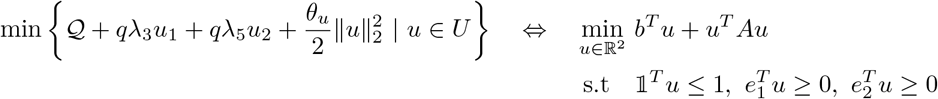

where 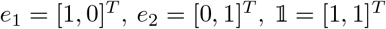, and

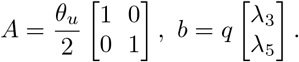

Let *u*^*^ be minimizer of above constrained QP problem, we observe that *u*^*^ has following closed form

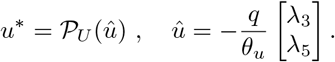

where 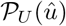 represents the projection of *û* onto *U*. Next, we derive the system of costate in Eq. (20). Note that 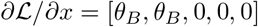, the terminal condition of costate equation is *λ*^*^(*t_f_*) = [*θ_f_, θ_f_*, 0, 0, 0]^*T*^ by Eq. (22). The Jacobian of the RHS of state equation is

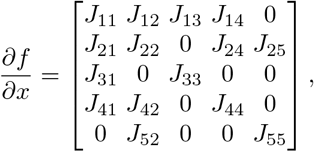

where

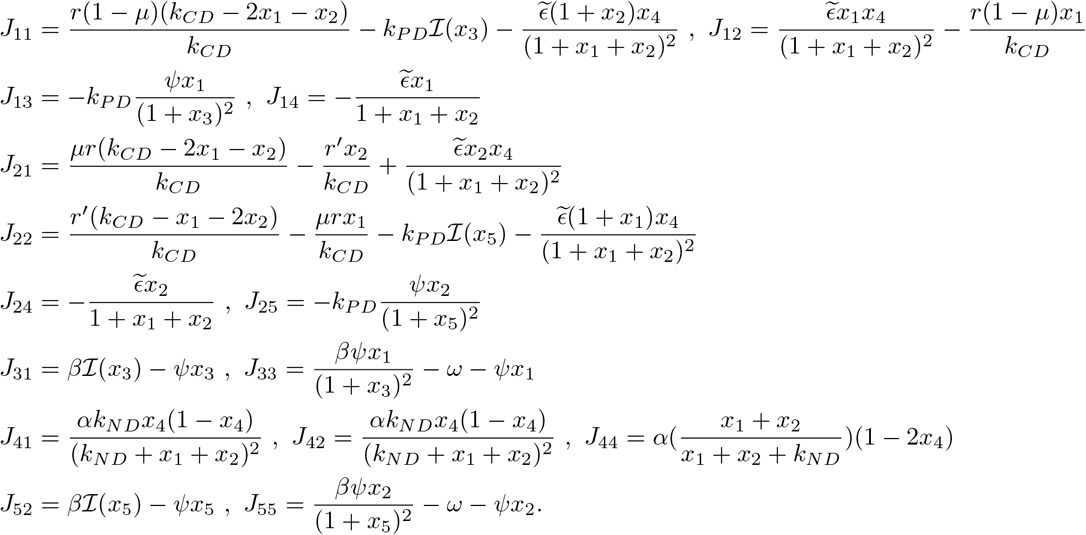

We have justified Eqs. (19)–(22).

### 3.3 Analysis of Optimal Control in Monophage Therapy

The monophage therapy can be modeled by a reduced form of system (1)–(5). The state vector of population densities is [*S, R, P_S_, I*]^*T*^, the corresponding population dynamics is modeled analogously to system (1)–(5) by excluding any terms associated with phage *P_R_*. Via the same scale transformations in Eq. (7) and Eq. (8), the scaled system of monophage therapy model is

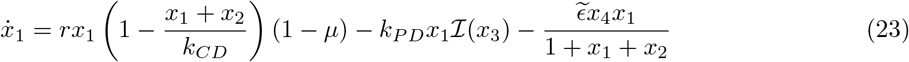

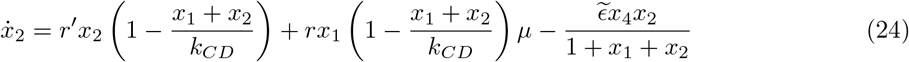

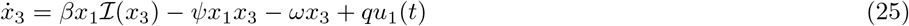

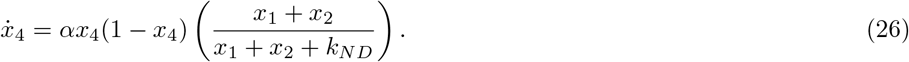

Note that the RHS of system (23)–(26) is the same as the RHS of system (9)–(13) by excluding all the terms associated with *x*_5_. Here, the space of controls is denoted by 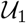, which is the set of Lebesgue-measurable functions *u*_1_: [*t*_0_, *t_f_*] → *U*_1_, where *U*_1_ = [0, 1]. The optimal control formulation is

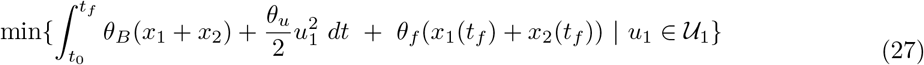

subject to the system (23)–(26).

The existence of optimal control is still guaranteed. The necessary conditions of optimal control in problem (27) are derived in Appendix B.

## 4 Numerical Results

### 4.1 The Hamiltonian-based Algorithm

We solve the optimal control problems by a Hamiltonian-based algorithm, this algorithm is presented with greater detail in [Hale et al., 2016, Wardi et al., 2016]. A salient feature is that the algorithm converges fast towards a near-optimal control (for a class of problems) if the Hamiltonian function is convex in *u* and can be minimized effectively and efficiently. Here, we briefly describe this algorithm. Two parameters are used to control the backtracking search, *η* ∈ (0, 1) and *s* ∈ (0, 1), and we used *η* = *s* = 0.5. At the *k^th^* iteration of the algorithm, *k* = 1,2,…, it starts with the *k^th^* control iteration *u^k^*, and computes from it the next iteration, *u*^*k*+1^, as follows.

1. Given a control input *u^k^*, compute the state trajectory x forward using numerical integration.
2. Compute the costate trajectory λ backward with terminal condition, the terminal condition.
3. For every *t* in a fine grid on the interval [*t*_0_, *t_f_*], compute *v*^*^(*t*) that minimizes the Hamiltonian. Interpolate the results by a zero-order hold (piecewise-constant interpolation) to define the control *v*^*^: = {*v*^*^(*t*): *t* ∈ [*t*_0_, *t_f_*]}. It serves as the steepest-feasible descent direction from *u^k^*.
4. Compute 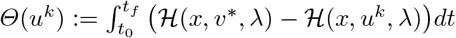.
5. Find 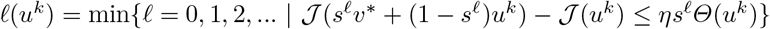.
6. Set 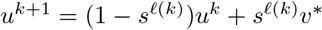.

The state trajectory *x*(*t*) and the costate trajectory λ(*t*) are numerically integrated by Euler’s method [Stoer and Bulirsch, 2013], the time step is *Δt* = 5 × 10^−4^. The convergence indicator |*Θ*(*u*)| measures the extent to which *u* fails to satisfy the PMP. The algorithm will be terminated either |*Θ*(*u*)| ≤ 10^−8^ or the maximum number of allowed iterations is reached.

### 4.2 Preliminaries of Simulations

#### Parameters

The model parameters and initial conditions of system (1)–(5) are given in Table 1. The *in silico* experiments run for 3 days post infection and all the treatments start at 2 hours after initialization (consistent with *in vivo* treatments in [Roach et al., 2017]), we thus set *t*_0_ = 2 hrs and *t_f_* = 72 hrs. We fix the regulator weights *θ_B_* = *θ_f_* = 1. We will tune the value of *θ_u_* to solve a practical variant of the original control problems (17) and (27).

**Table 1.** Parameters and initial conditions in the system (1)–(5)

| Parameters | Value | Source |
| --- | --- | --- |
| $r$, maximum growth rate of bacteria | $0.75 \text{ h}^{-1}$ | [Roach et al., 2017] |
| $r'$, maximum growth rate of phage-resistant bacteria | $0.675 \text{ h}^{-1}$ | [Roach et al., 2017] |
| $\mu$, probability of emergence of phage-resistant mutant per cellular division | $2.85 \times 10^{-8}$ | [Roach et al., 2017] |
| $K_C$, carrying capacity of bacteria (wild-type) | $1.0 \times 10^{10} \text{ CFU/g}$ | [Roach et al., 2017] |
| $K_C$, carrying capacity of bacteria (immunodeficient hosts) | $8.5 \times 10^{11} \text{ CFU/g}$ | [Roach et al., 2017] |
| $\beta$, burst size of phage | 100 | [Roach et al., 2017] |
| $w$, decay rate of phage | $0.07 \text{ h}^{-1}$ | [Roach et al., 2017] |
| $\epsilon$, killing rate parameter of immune response | $8.2 \times 10^{-8} \text{ g/(h cell)}$ | [Roach et al., 2017] |
| $\alpha$, maximum growth rate of immune response | $0.97 \text{ h}^{-1}$ | [Roach et al., 2017] |
| $K_I$, maximum capacity of immune response | $2.4 \times 10^7 \text{ cell/g}$ | [Roach et al., 2017] |
| $K_I$, maximum capacity of immune response (no innate immune activation) | same as $I_0$ | [Roach et al., 2017] |
| $K_D$, bacterial concentration at which immune response is half as effective | $4.1 \times 10^7 \text{ CFU/g}$ | [Roach et al., 2017] |
| $K_N$, bacterial concentration when immune response growth rate is half its maximum | $1.0 \times 10^7 \text{ CFU/g}$ | [Roach et al., 2017] |
| $\phi$, adsorption rate of phage | $5.4 \times 10^{-8} \text{ g/(h PFU)}$ | [Roach et al., 2017] |
| $P_C$, phage density at half saturation | $1.5 \times 10^7 \text{ PFU/g}$ | [Roach et al., 2017] |
| $S_0$, initial bacterial density | $7.4 \times 10^7 \text{ CFU/g}$ | [Roach et al., 2017] |
| $S_0$, initial bacterial density (immunodeficient hosts) | $7.4 \times 10^5 \text{ CFU/g}$ | [Roach et al., 2017] |
| $R_0$, initial phage-resistant bacterial density | $1 \text{ CFU/g}$ | [Roach et al., 2017] |
| $I_0$, initial immune response | $2.7 \times 10^6 \text{ cell/g}$ | [Roach et al., 2017] |
| $I_0$, initial immune response (immunodeficient hosts) | $3 \times 10^6 \sim 8.5 \times 10^6, \text{ cell/g}$ | this study |
| $P_S(0)$, initial WT phage density | $0 \text{ PFU/g}$ | this study |
| $P_R(0)$, initial host-range mutant phage density | $0 \text{ PFU/g}$ | this study |
| $\rho_{max}$, maximal phage dose injection rate | $10^9 \text{ PFU/(h g)}$ | this study |

#### Practical Treatment Objective

The goal of the optimal control framework described in Sect. 2.2 is to minimize the total bacterial population and penalizing the treatment costs by optimizing the scaled phage injection rates. However, from a practical therapeutic perspective, we may want to find the minimal phage dosage required to eliminate bacteria instead. Since the differential equation system (1)–(5) is continuous, we assume the bacterial population is eliminated if there exists a time *τ* ∈ [0, *t_f_*] such that *S*(*τ*) + *R*(*τ*) ≤ *n*_ext_, where *n*_ext_ = 1 CFU/g is the hard threshold of bacteria elimination.

The objective of practical therapy can thus be formalized as a constrained optimal control problem that minimizes the integral of phage injection rates subject to the constraint of bacteria elimination in a *hybrid system* where the state equations are different before and after bacterial elimination. The discontinuous nature of bacterial elimination events poses considerable numerical challenge in explicitly solving the constrained hybrid optimal control problem. As a result, we utilize the ordinary control formulations (17) and (27) to achieve the goal of eliminating bacteria with minimum dosage via a heuristic approach. In this approach, we adjust the regulator weight *θ_u_* and locate the highest value that results in bacterial elimination. The total phage dosage is then computed to find the minimum dosage corresponding to bacterial elimination.

#### Minimal Phage Dosage and Regulator Weight θ_u_

Here, we detail the procedure of achieving the practical control objective by tuning regulator weight *θ_u_* in a certain range. We search *θ_u_* in the interval [10^−11^, 10^11^], ranging from negligible treatment costs (*θ_u_* = 10^−11^) to dominating treatment costs (*θ_u_* = 10^11^) in the control objective. For any fixed *θ_u_* ∈ [10^−11^, 10^11^], we can numerically solve the optimal control problems (17) and (27) via a Hamiltonian-based algorithm. It is also easy to check if the optimal control solution effectively eliminates bacterial populations based on the artificial threshold we introduced.

In order to find the treatment (*i.e*., profiles of phage injection rate) that can eliminate bacteria with minimal dosage, we sweep over the range of *θ_u_* and extract all the ‘*θ_u_*’ values that lead to successful bacterial elimination. Assuming such ‘*θ_u_*’ values exist, we then find the minimum effective phage dosage by computing the integral of phage injection rate that corresponds to the highest ‘*θ_u_*’ value resulting in bacterial elimination, e.g., dosage of a treatment that has two types of phage is 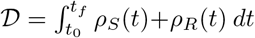. In the case that no optimal control treatment can eliminate the bacteria with *θ_u_* in the range of [10^−11^, 10^11^], we assume phage therapy would fail under all reasonable phage dosages in such conditions.

However, the computational costs of finding the value of *θ_u_* corresponding to the minimum effective dosage 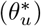 via brute-force searching are very high. Intuitively, we note that if bacteria is eliminated for a given 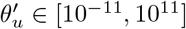, then bacteria would also be eliminated for all 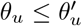, i.e., relaxing penalization on treatment costs would always eliminate bacteria. Likewise, if bacteria is not eliminated for 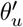, then bacteria is also not eliminated for all 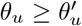 as higher penalization on treatment costs would lead to a less effective treatment.

Building upon these intuitions, we implement a binary search algorithm to find 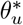. We first evaluate the two boundary cases (*θ_u_* = 10^−11^ and *θ_u_* = 10^11^) and assume that phage therapy would fail in general if the optimal treatment corresponding to *θ_u_* = 10^−11^ fails. In addition, if the optimal treatment corresponding to *θ_u_* = 10^11^ can successfully eliminate bacteria, its associated phage dosage is identified as the minimum effective dosage. If both of the aforementioned boundary conditions are not satisfied, we compute the optimal treatment corresponding to an intermediate weight *θ_ui_* = 10^(*L+R*)/2^ where *L* = −11 and *R* = 11 for the two boundaries. If optimal treatment with *θ_ui_* works, we update left searching boundary *L* ← (*L* + *R*)/2; otherwise, we update right searching boundary *R* ← (*L* + *R*)/2. We iterate this procedure for *n* = 8 times (corresponding to a precision of about 22/2^8^ ≈ 0.08 at power scale, where 22 is the length of power scale range for *θ_u_* ∈ [10^−11^, 10^11^]) to estimate the value of 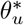.

#### Classification of Injection Strategies

In this study, we focus on three types of injection strategies: an optimal control strategy in monophage therapy, namely, one dimensional optimal control (*1D-OC*); an optimal control strategy using multiple types of phage, namely, two dimensional optimal control (*2D-OC*); and a *practical therapeutic treatment* using either single or multiple types of phage. The 1D-OC and 2D-OC are optimal controls solved numerically from problems (27) and (17) respectively. However, 1D-OC and 2D-OC are usually continuous signals, which cannot be directly implemented in (current) clinical treatment, we thus have to convert the continuous treatment to a (discrete) multi-dose treatment.

In this paper, we only focus on developing multi-dose treatment guided by 2D-OC treatment in immunodeficient scenarios. There are two types of phage in 2D-OC and we allow one-time dose injection for each type of phage in the practical therapeutic treatment. The timings of injecting phage *P_S_* and phage *P_R_* (in practical therapeutic treatment guided by 2D-OC) are denoted by *T_P_S__* and *T_P_R__* respectively. We define *T_P_S__* = *min*{*τ* ∈ [*t*_0_, *t_f_*] |*ρ_S_*(*τ*) ≥ *ρ_S_*(*t*) ∀*t* ∈ [*t*_0_, *t_f_*]}, *i.e*., the first time that injection rate of phage *P_S_* arrives its maximal rate. *T_P_R__* is defined in the same way as *T_P_S__*. We define the integral of phage injection rate *ρ_S_*(*t*) over [*t*_0_,*t_f_*] as 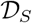, and the analogous integral ofthe second phage type as 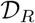. In doing so, given a 2D-OC treatment (*ρ_S_*(*t*), *ρ_R_*(*t*)), the practical therapeutic treatment guided by this 2D-OC treatment is: injecting 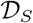 amount of phage *P_S_* at time *T_P_S__* and injecting 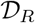 amount of phage *P_R_* at time *T_P_R__*. The *in silico* experiments using practical therapeutic treatment assumes bacteria and phage are eliminated locally when their population densities drop below a threshold of 1 *g*^−1^. As the practical therapeutic treatment guided by 2D-OC is only an approximation of the optimal treatment, it is possible for the practical strategy to fail to eliminate bacteria. In this situation, we iteratively amplify the dosages (ten percent higher for each step) of the two phage types used in practical therapeutic treatment while fixing the timings of phage injection until bacterial populations are eliminated.

### 4.3 Monophage Therapy in Immunocompetent Hosts

For immunocompetent hosts with intact immune activation, monophage therapy can be highly effective in curing bacterial infections [Roach et al., 2017]. However, phage therapy can still fail when the phage decay rate is high, or when the phage infection rate is low due to inefficient phage strains or partial resistance [Leung and Weitz, 2017, Roach et al., 2017]. To explore these potential modes of failure, we set the phage infection rate to be *ϕ* = 3.38 × 10^−8^ g/(h PFU), slightly lower than the estimated value *ϕ* = 5.4 × 10^−8^ g/(h PFU) in [Roach et al., 2017]. We then plot the performances of 1D-OC strategy by their minimal phage dosages for eliminating the bacterial population given a range of phage decay rate *ω* ∈ [10^−2^, 10^2^](*h*^−1^).

In Fig. 2, we find that the minimum phage dosage needed by 1D-OC strategies to eliminate the bacteria increases monotonically with the phage decay rate. In addition, the dosage increase becomes extremely rapid at decay rate *ω* ∼ 1 h^−1^ and 1D-OC therapy fails for all practical dosages when *ω* > 2.5 h^−1^. The time series of population dynamics at slow phage decay rate are plotted in Fig. 3A. The single-impulse like optimal injection rate shows that the 1D-OC treatment (in this case) is approximately a single-dose treatment: injecting a small amount of phage (about 5 × 10^2^ PFU) at the very beginning of treatment 2 hours post infection. In doing so, the optimized treatment reduces the phage-sensitive bacterial population quickly and controls the emergence of resistant bacteria in an early stage. On the other hand, when phage decay at a fast rate, the 1D-OC strategy maintains the phage concentration and treatment efficacy by continuously injecting phage into the system over a longer period of time (see Fig. 3B).

**Fig. 2.**
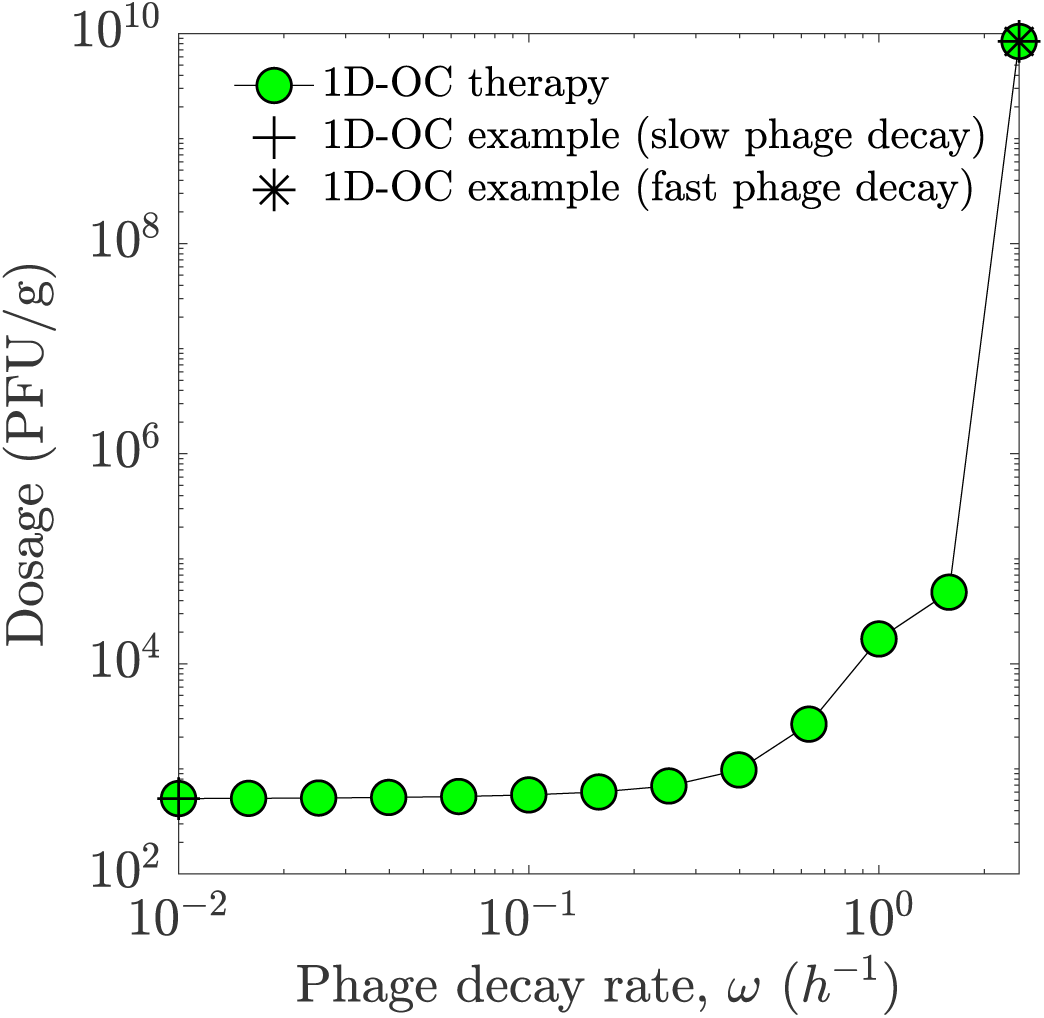
Minimal phage amount for eliminating bacterial cells using 1D-OC strategy (green dots). There is no 1D-OC treatment that can eliminate bacteria in the regime of high phage decay rate (*ω* ≥ 2.5h^−1^). Two 1D-OC examples are provided in Fig. 3: the corresponding time series of population densities and injection rate trajectories. See model parameters and simulation details in Sect. 4.2.

**Fig. 3.**
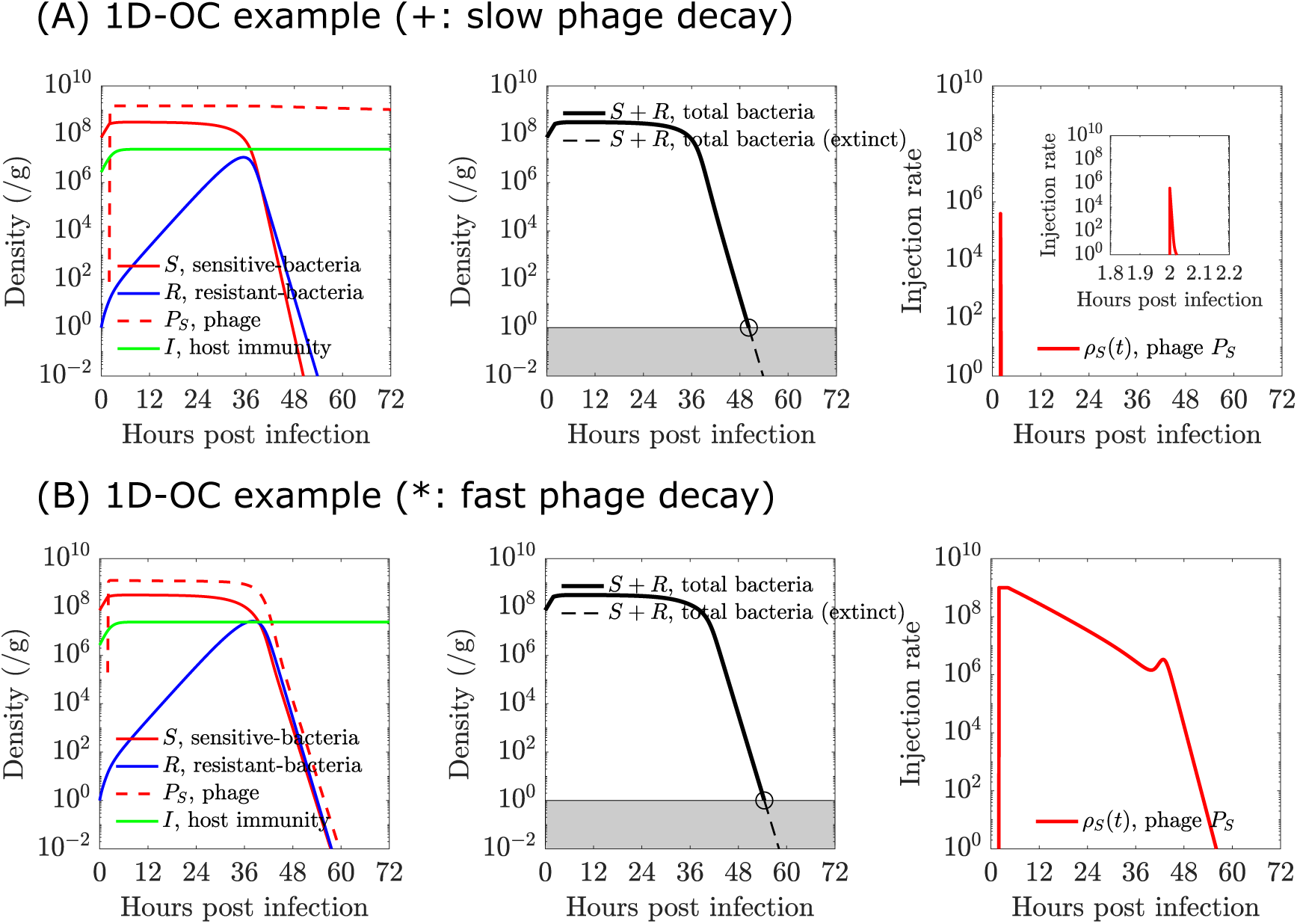
Time series of population densities and optimal injection rates of two 1D-OC examples (labeled in Fig. 2): slow phage decay (*ω* = 0.01h^−1^) and fast phage decay (*ω* = 2.5h^−1^). (A) Optimal injection rate, *ρ_S_* (*t*), is obtained by solving control problem (27) with tuned regulator weight *θ_u_* = 10^11^. Bacteria is eliminated around 50 hrs post infection. (B) Optimal injection rate, *ρ_S_* (*t*), is obtained by solving control problem (27) with tuned regulator weight *θ_u_* = 10. Bacteria is eliminated around 50 hrs post infection. See model parameters and simulation details in Sect. 4.2.

### 4.4 Phage Therapy in Immunodeficient Hosts

Previous work has shown that a deficient immune response may lead to failure of phage therapy in an acute pneumonia system [Roach et al., 2017]. Here, we explore whether phage combination therapy (phage cocktails) that includes a host-range mutant phage targeting resistant bacteria can restore therapeutic effectiveness in immuno-deficient hosts, and identify optimal ways to achieve that. In the immunodeficient model, immune signaling is assumed to be absent such that the immune response intensity is maintained at a basal level, *I* = *I*_0_. In addition, the initial density of sensitive bacteria that is 2 log lower than the initial density of sensitive bacteria because immunodeficient hosts are highly susceptible to bacterial infection [Roach et al., 2017]. We further assume that the wild-type and host-range mutant phage target the phage-sensitive and phage-resistant bacteria respectively with no cross infectivity. First, we compute 2D-OC treatments corresponding to minimal phage dosages given a range of basal-level immune density. Then, we test a practical approximation of the 2D-OC treatment by converting the 2D-OC phage injection profile into discrete doses of each phage strain (see Sect. 4.2 for details). For each basal-level immune density, we evaluate the performance of the practical therapeutic treatment guided by the 2D-OC treatment at the same basal-level immune density.

The numerical results show that the the minimal phage amount of both 2D-OC and practical therapies for eliminating bacterial cells decreases with increasing basal-level immune density (see Fig. 4A). Moreover, for practical therapeutic treatments in the entire range of the basal-level immune density, the optimal timings of injecting the two types of phage are both at the very beginning of treatment (*i.e*., *T_P_S__* = *T_P_R__* ≈ 2 hrs). Furthermore, given our model and parameter assumptions the optimal control algorithm always result in a dosage of phage *P_S_* about ten times higher than the dosage of phage *P_R_* (*i.e*., 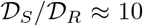), see Fig. 4B. When the basal-level immune density is very high, *I*_0_ ≈ 8.5 × 10^6^ (cell/g), the practical treatment only needs a small amount of phage *P_S_* (about 10^2^ PFU) to eliminate bacteria (see Fig. 4B and Fig. S1 in Appendix C.). To investigate the phage delivery schedules identified by the optimal control strategies and understand the advantages of phage combination therapy, we compare the injection rate and population dynamics of 1D-OC, 2D-OC and practical therapeutic strategies in Figs. 5 and 6.

**Fig. 4.**
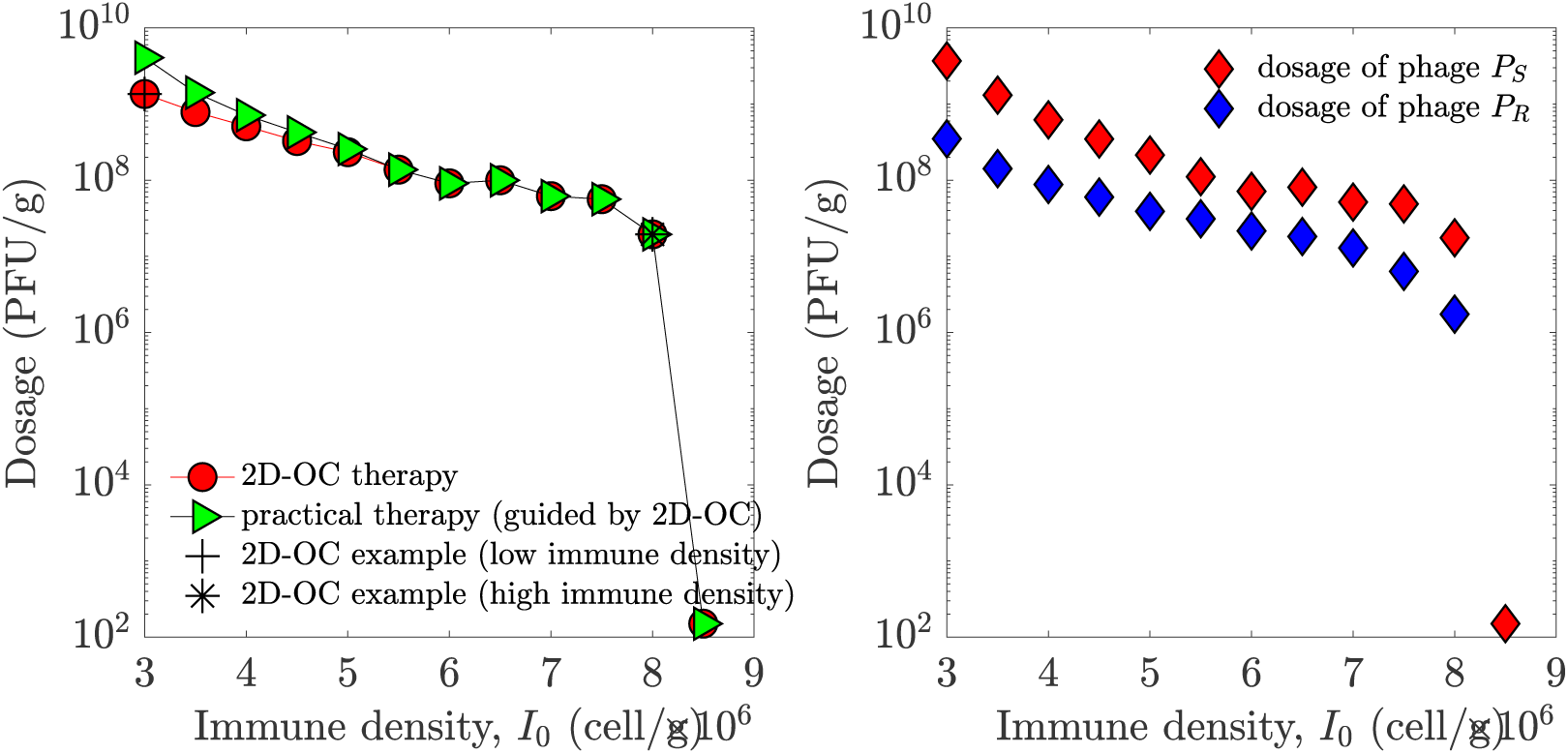
Minimal phage amount for eliminating bacterial cells using 2D-OC strategy (left) and the practical therapeutic treatment (right). (Left) Minimal phage amount for eliminating bacterial cells using 2D-OC strategy (red) and practical therapeutic treatment guided by 2D-OC strategy (green). Two in silico experiments (high immune density *I*_0_ = 8×10^6^ cell/g and low immune density *I*_0_ = 3 × 10^6^ cell/g) are provided in Figs. 5 and 6 respectively. (Right) Dosage of phage *P_S_* (red) and dosage of phage *P_R_* (blue) used in practical therapeutic treatment. The timings of injecting two types of phage dose are both about two hours post infection (i.e., *T_P_S__* = *T_P_R__* ≈ 2 hrs). Note that the total dosage of two types of phage (add up of *P_S_* phage dosage and *P_R_* phage dosage) is the green curve in the left panel. See model parameters and simulation details in Sect. 4.2.

**Fig. 5.**
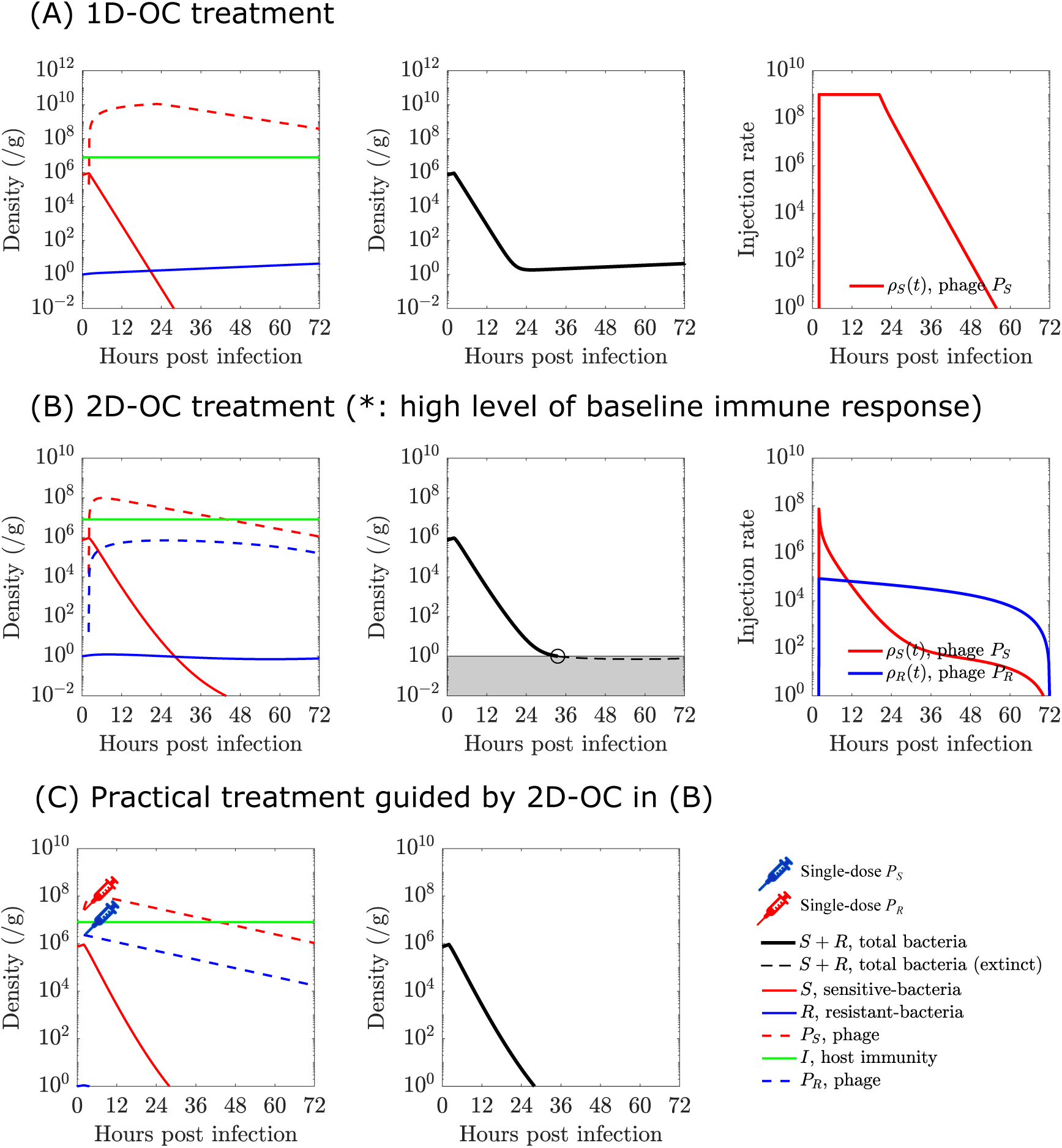
Comparison of time series of population densities with different treatments in the high level of baseline immune response, *I*_0_ = 8 × 10^6^ cell/g. (A) Optimal injection rate, *ρ_S_*(*t*), is obtained by solving control problem (27) with tuned regulator weight *θ_u_* = 10^−11^ (the smallest regulator weight on treatment costs). There does not exist curative 1D-OC treatment due to the outgrowth of phage-resistant bacteria *R*. (B) Optimal injection rate, *ρ_S_*(*t*) and *ρ_R_*(*t*), is obtained by solving control problem (17) with tuned regulator weight *θ_u_* = 10. Bacteria is eliminated around 30 hrs post infection. (C) The practical therapeutic treatment is obtained from optimal injection rate in (B): two doses, *P_S_* phage dose and *P_R_* phage dose, both are injected at two hours post infection with amount of 2.6 × 10^7^ PFU and 2.3 × 10 ^6^ PFU respectively. Bacteria is eliminated around 30 hrs post infection. See model parameters and simulation details in Sect. 4.2.

**Fig. 6.**
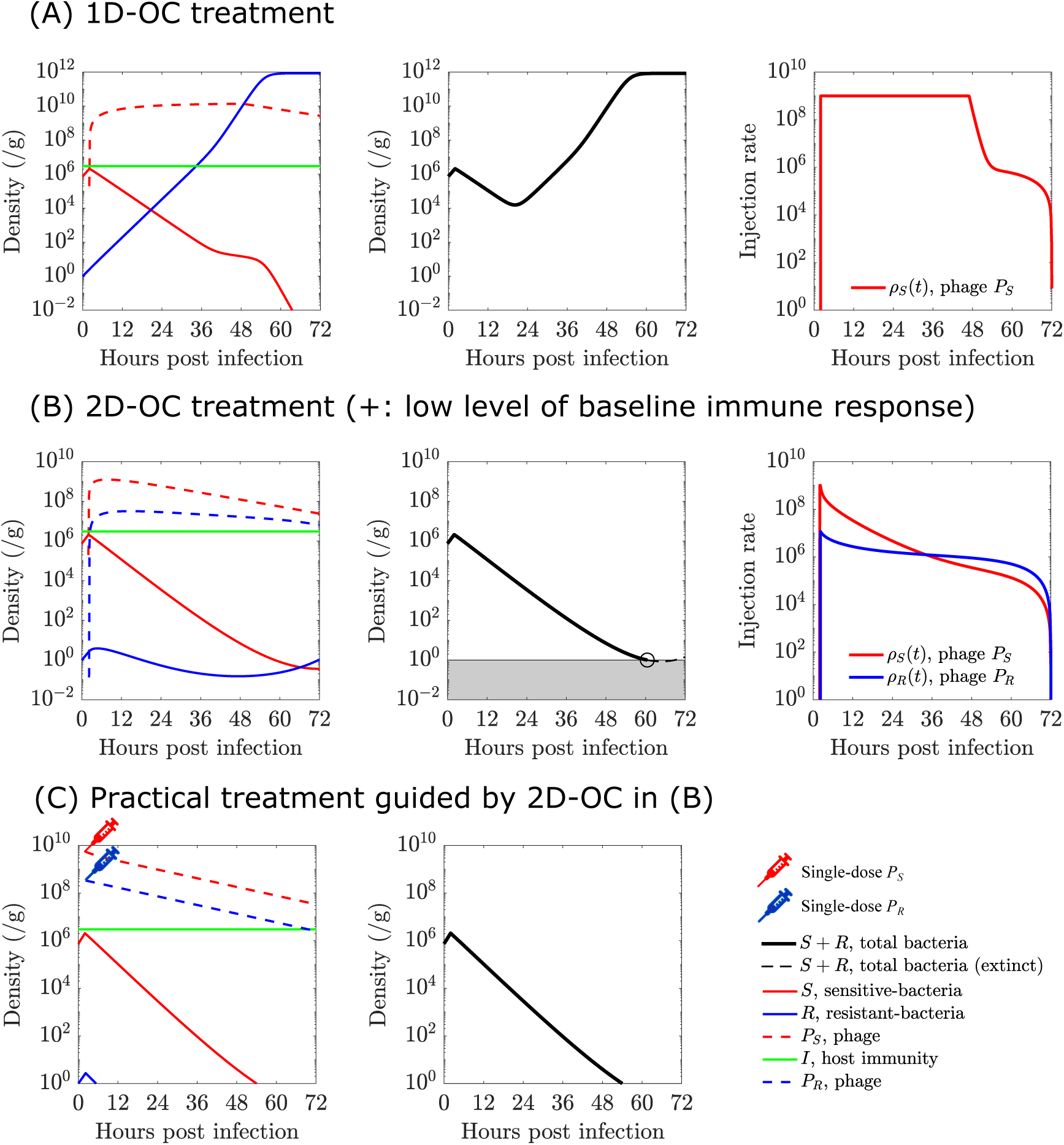
Comparison of time series of population densities with different treatments in the low level of baseline immune response, *I*_0_ = 3 × 10^6^ cell/g. (A) Optimal injection rate, *ρ_S_*(*t*), is obtained by solving control problem (27) with tuned regulator weight *θ_u_* = 10^−11^ (the smallest regulator weight on treatment costs). There does not exist curative 1D-OC treatment due to the outgrowth of phage-resistant bacteria R. (B) Optimal injection rate, *ρ_S_*(*t*) and *ρ_R_*(*t*), is obtained by solving control problem (17) with tuned regulator weight *θ_u_* = 10^−2^. Bacteria is eliminated around 60 hrs post infection. (C) The practical therapeutic treatment is obtained from optimal injection rate in (B): two doses, *P_S_* phage dose and *P_R_* phage dose, both are injected at two hours post infection with amount of 3 × 10^9^ PFU and 3 × 10^8^ PFU respectively. Bacteria is eliminated around 60 hrs post infection. See model parameters and simulation details in Sect. 4.2.

For the 1D-OC case at high level of immune response (see Fig. 5A), the optimal phage dosing occurs over a time period of nearly 60 hours with injection rate starting at around the maximal value of 10^9^ PFU/(h g) and decreasing after 24 hours. The immunodeficiency necessitated a much higher dosage of phage sustained over a long period of time even for a low initial bacterial inoculum. The aggressive phage dosing results in a roughly exponential decrease in phage-sensitive bacteria, but the phage-resistant bacteria is not effectively controlled and increases slightly in population. Phage-resistant bacteria is even more problematic at low baseline immune level (see Fig. 6A), where the resistant mutants grow exponentially until they reach the carrying capacity despite a more aggressive phage dosing that maintains a maximum injection rate until 48 hours post infection. The failure of phage therapy in these cases is a result of the deficient immune response not being able to control phage-resistant bacteria, as confirmed by the population dynamics at sufficiently high baseline immune response (see Fig. S1 in Appendix C.) which shows effective control of both phage-sensitive and phage-resistant bacterial populations at much lower phage dose.

Figs. 5B-C and 6B-C show how judicious use of a phage cocktail in the 2D-OC strategy can improve the robustness of therapeutic success. In Fig. 5B, the host-range mutant phage is injected for the entire dosing schedule to preemptively inhibit the phage-resistant bacteria. Initially, wild-type phage targeting the phage-sensitive bacteria is injected at a higher dose than the host-range mutant phage as the initial inoculum consists mostly of sensitive bacteria. That balance is shifted as the relative proportion of resistant bacteria increases and the host-range mutant phage becomes the major component of the cocktail at around 12 hrs post infection. Thus, the optimal injection rates (in 2D-OC treatment) exhibit interesting and complex temporal patterns. However, in real clinical treatments, it is not (yet) feasible to implement such a complicated phage delivery schedules. In Fig. 5C, the practical therapeutic treatment, i.e., a discretized form of the continuous 2D-OC treatment, also shows efficacy in eliminating the bacteria. In the case of low baseline immune response, the temporal patterns in treatment strategies (2D-OC and practical treatments) are similar to the case of high baseline immune response (see Figs. 6B-C and compare to Figs. 5B-C), but the dosage of two types of phage used in treatments are much higher. This is consistent with our previous work showing that the host immune status may be a critical factor determining phage therapy efficacy [Roach et al., 2017]. In addition, for low immune level the time of switching over to a strategy dominated by the counter-resistant phage is delayed to about 36 hours after infection as opposed to around 12 hours in the high immune baseline case.

## 5 Discussion

In this paper, we developed a control-theoretic framework to optimize the use of monophage treatment and phage cocktails for treating bacterial infections in immunodeficient hosts or in other scenarios such as high phage decay rate where standard phage therapy is likely to fail. By incorporating phage administration as the control in a mathematical model describing the nonlinear interactions between phage, pathogenic bacteria and host immunity, we derive a Hamiltonian-based algorithm to numerically minimize bacterial burden while limiting the phage dose. Our results indicate that optimal control may provide important insights to guide clinical use of phage therapy. In particular, a single dose of phage may be sufficient to treat immunocompetent patients when the phage clearance rate is low, whereas phage administration may need to be sustained over a longer period when phage clearance is fast. In immunodeficient hosts, our results suggest that the success of optimal administration of phage cocktails can largely be reproduced in a simplified, discretized version of the optimal therapy that would be easier to implement practically. The only trade-off observed for the simplified practical therapy is a slight increase of the minimum effective dose in cases of severe immunodeficiency. Our optimal control framework indicates that a single phage strain may be effective for therapy at relatively high immunity levels, but the use of multiple therapeutic phage is required for low immune intensities.

To ensure that the optimal control problem remains mathematically tractable, a number of simplifying assumptions have been made. For example, within-host dynamics is assumed to be deterministic, whereas biological processes such as mutations are inherently stochastic. The gap between mathematical models and complex clinical trials can be even wider due to various confounding factors. In addition, the optimal control solutions are solved based on specific objective functional and model parameters. Hence, different sets of parameters, models, and the objective costs could yield different suggested treatments such that the robustness of our optimal treatments may not be guaranteed. Our model also focuses on acute infections, and did not consider spatial heterogeneity or cocktails consisting of more than two phage strains. These issues could be addressed in future work by extending our modeling framework to incorporate stochastic control [Hashemian and Armaou, 2017, Fleming and Rishel, 1976] in spatial models with multiple strains of phage. In doing so, it will be important to consider host adaptive immunity, which is important in chronic infections and can generate specific responses against phage [Hodyra-Stefaniak et al., 2015, Gorski et al., 2012]. Finally, we note that host-range phage mutants may be able to infect multiple strain types [Flores et al., 2011]; hence future work should also address how to optimally combine phage with overlapping host ranges.

In conclusion, the theoretical framework presented in this paper is intended to help advance the rational design of monophage and phage cocktail therapy. Phage cocktails have been proposed as a solution to tackle phage resistance and broaden the antimicrobial spectrum of phage preparations [Chan et al., 2013, Tanji et al., 2004, Zhang et al., 2010], but it is often unclear how to optimize their composition to obtain maximum effectiveness. Part of the difficulty in determining the appropriate dosage and composition of phage treatments is due to the self-amplifying nature of phage. Our work demonstrates how control theory can be applied to optimize the dose and timing of therapeutic agents that have the ability to proliferate *in vivo*, and provide insights, both of a conceptual and practical nature, in the development of phage therapy. In doing so, our framework can also be extended to other therapeutic contexts with replicating agents, such as the use of probiotic bacteria [Manzanares et al., 2016, Parker et al., 2018], as well as cancer therapies involving oncolytic viruses [Lawler et al., 2017, Fukuhara et al., 2016] and live immune cells [June et al., 2018, Bol et al., 2016].

## Appendix

## A. Implementation of Projection Operator 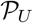

Here we present a closed form of projection operator 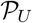 via a geometric approach, recall that 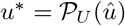 in Theorem 1, then we have following:

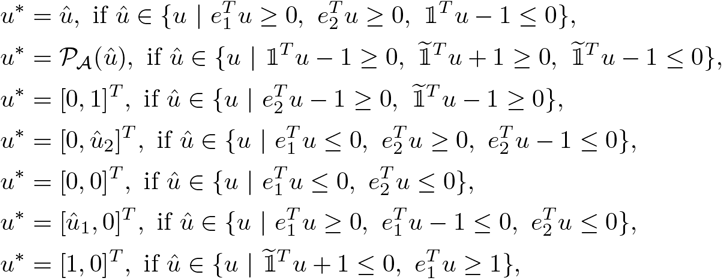

where 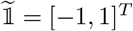 and 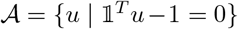. Computing the projection of *û* on to 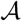 is straightforward. The orthogonality principle yields 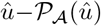 must be colinear with 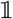, also 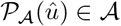, *i.e*., 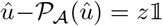 and 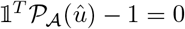. This yields 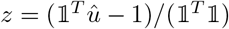 and thus 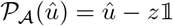.

## B. The Optimality System of Optimal Control in Monophage Therapy

Here, we derive the necessary conditions for the optimal control problem (27) via PMP.

### Theorem 2

*If* 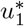 *is an optimal control solves problem (27), and x*^*^(*t*) *and* λ^*^(*t*) *are the corresponding state trajectory and costate trajectory, then*

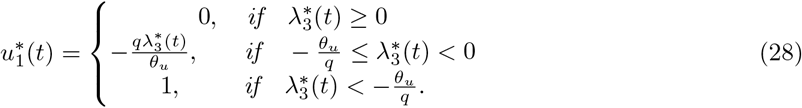

*Proof*. According to PMP, if 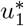 is an optimal control of problem (27), and if *x*^*^(*t*) and λ^*^(*t*) are the corresponding state trajectory and costate trajectory, then the following equations are satisfied,

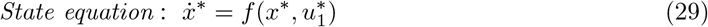

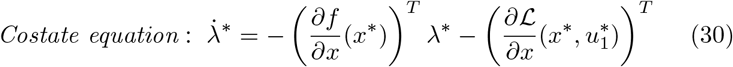

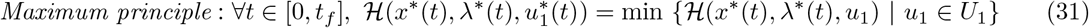

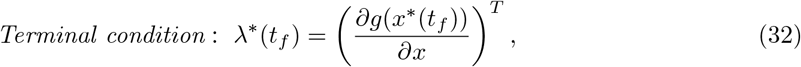

where 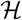 is the Hamiltonian. Define the Hamiltonian as 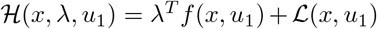, we find that 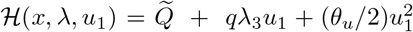, where 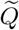 is the collection of terms that has no argument in *u*_1_. Minimizing 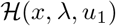 over *u*_1_ ∈ *U*_1_ yields Eq. (28). The costate equation with terminal condition is

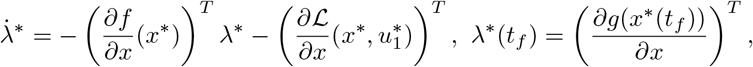

where 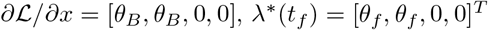. The Jacobian is

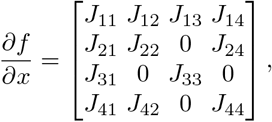

where

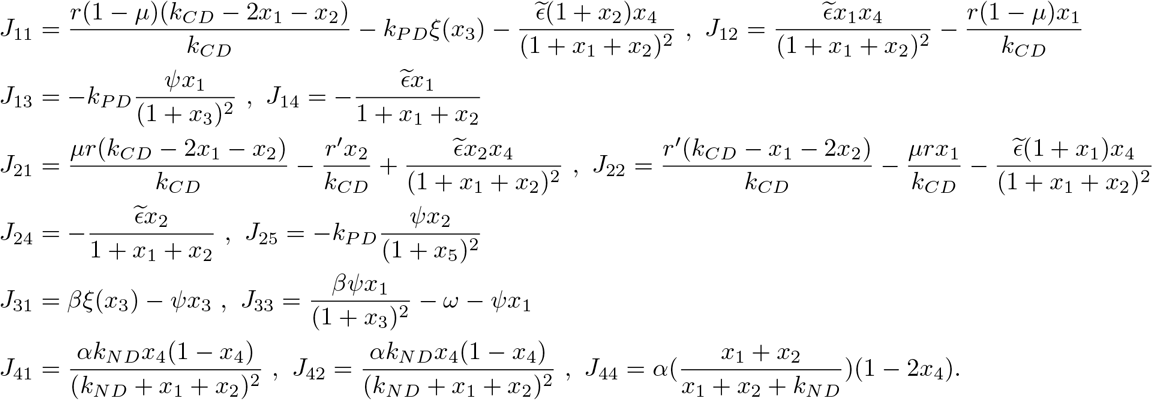

We have justified Eqs. (29)-(32).

**Fig. S1.**
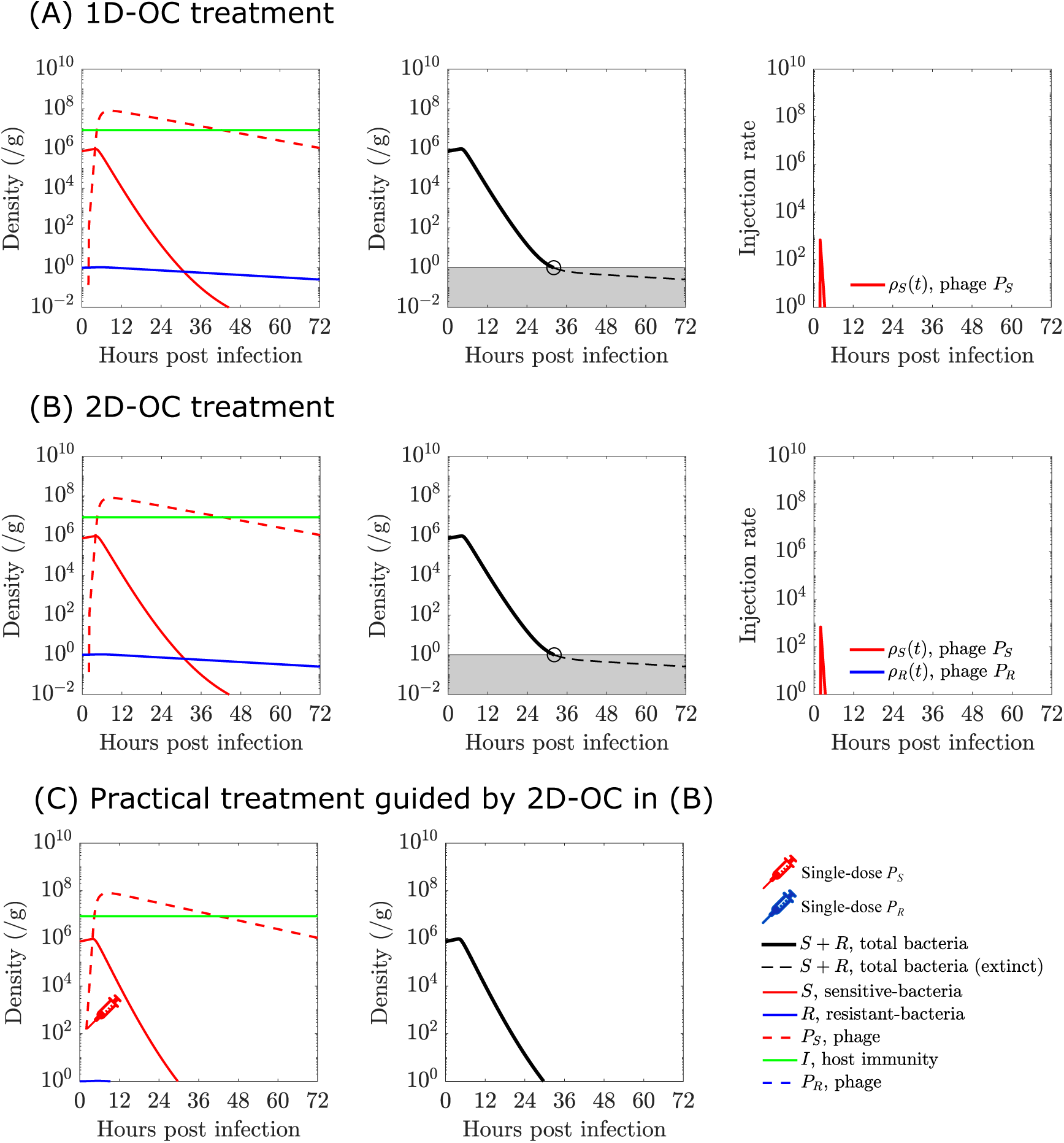
Comparison of time series of population densities with different treatments in the high level of baseline immune response, *I*_0_ = 8.5 × 10^6^ cell/g. (A) Optimal injection rate, *ρ_S_* (*t*), is obtained by solving control problem (27) with tuned regulator weight *θ_u_* = 10^11^ (the largest regulator weight on treatment costs). Bacteria is eliminated around 30 hrs post infection. (B) Optimal injection rate, *ρ_S_* (*t*) and *ρ_R_*(*t*), is obtained by solving control problem (17) with tuned regulator weight *θ_u_* = 10^11^. Note that the optimal injection rate of phage *P_R_* is nearly zero, *i.e*., *ρ_R_*(*t*) ≈ 0 ∀*t* ∈ [*t*_0_, *t_f_*]. Thus, the optimal injection rates solved from 2D-OC and 1D-OC are nearly identical, *i.e*., *ρ_S_*(*t*) is a single-pulse signal centered at t=2 hrs. Bacteria is eliminated around 30 hrs post infection. (C) The practical therapeutic treatment is obtained from optimal injection rate in (B): single-dose, *P_S_* phage dose, is injected at two hours post infection with amount of 5× 10^2^PFU. Bacteria is eliminated around 30 hrs post infection. See model parameters and simulation details in Sect. 4.2.

## C. Effective Single-Dose Treatment in Immunodeficient Hosts (Baseline Immune Response is Sufficiently High)

When the baseline immune response is sufficiently high in immunodeficient hosts, all the treatment strategies (1D-OC, 2D-OC and practical treatments) can eliminate bacteria with a low dose of phage *P_S_* injected at very beginning of treatment (see Fig. S1 in Appendix C.).

## Acknowledgements

The work was supported by a grant from the Army Research Office W911NF-14-1-0402 (to JSW) and a grant from the National Institutes of Health 1R01AI146592-01 (to JSW and LD).

## References

Edoardo Beretta and Yang Kuang. Modeling and analysis of a marine bacteriophage infection. Mathematical Biosciences, 149(1):57–76, 1998.

Kbenesh Blayneh, Yanzhao Cao, and Hee-Dae Kwon. Optimal control of vector-borne diseases: treatment and prevention. Discrete and Continuous Dynamical Systems B, 11(3):587–611, 2009.

Kalijn F Bol, Gerty Schreibelt, Winald R Gerritsen, I Jolanda M De Vries, and Carl G Figdor. Dendritic cell–based immunotherapy: state of the art and beyond, 2016.

ET Camacho, LA Melara, MC Villalobos, and Stephen Wirkus. Optimal control in the treatment of retinitis pigmentosa. Bulletin of mathematical biology, 76(2):292–313, 2014.

Filippo Castiglione and Benedetto Piccoli. Optimal control in a model of dendritic cell transfection cancer immunotherapy. Bulletin of Mathematical Biology, 68(2):255–274, 2006.

Benjamin K Chan, Stephen T Abedon, and Catherine Loc-Carrillo. Phage cocktails and the future of phage therapy. Future microbiology, 8(6):769–783, 2013.

Benjamin K Chan, Mark Sistrom, John E Wertz, Kaitlyn E Kortright, Deepak Narayan, and Paul E Turner. Phage selection restores antibiotic sensitivity in mdr pseudomonas aeruginosa. Scientific reports, 6:26717, 2016.

Benjamin K Chan, Paul E Turner, Samuel Kim, Hamid R Mojibian, John A Elefteriades, and Deepak Narayan. Phage treatment of an aortic graft infected with pseudomonas aeruginosa. Evolution, medicine, and public health, 2018(1):60–66, 2018.

Ana-Maria Croicu. Short-and long-term optimal control of a mathematical model for HIV infection of CD4^+^T cells. Bulletin of mathematical biology, 77(11):2035–2071, 2015.

Ana-Maria Croicu. An optimal control model to reduce and eradicate anthrax disease in herbivorous animals. Bulletin of Mathematical Biology, 81(1):235–255, 2019.

Ana-Maria Croicu, Angela M Jarrett, NG Cogan, and M Yousuff Hussaini. Short-term antiretroviral treatment recommendations based on sensitivity analysis of a mathematical model for hiv infection of cd4+ t cells. Bulletin of mathematical biology, 79(11):2649–2671, 2017.

Rebecca V Culshaw, Shigui Ruan, and Raymond J Spiteri. Optimal hiv treatment by maximising immune response. Journal of Mathematical Biology, 48(5):545–562, 2004.

Lisette G de Pillis, K Renee Fister, Weiqing Gu, Tiffany Head, Kenny Maples, Todd Neal, Anand Murugan, and Kenji Kozai. Optimal control of mixed immunotherapy and chemotherapy of tumors. Journal of Biological systems, 16(01):51–80, 2008.

Rebekah M Dedrick, Carlos A Guerrero-Bustamante, Rebecca A Garlena, Daniel A Russell, Katrina Ford, Kathryn Harris, Kimberly C Gilmour, James Soothill, Deborah Jacobs-Sera, Robert T Schooley, et al. Engineered bacteriophages for treatment of a patient with a disseminated drug-resistant mycobacterium abscessus. Nature medicine, 25(5):730, 2019.

Nicolas Dufour, Raphaëlle Delattre, Anne Chevallereau, Jean-Damien Ricard, and Laurent Debarbieux. Phage therapy of pneumonia is not associated with an overstimulation of the inflammatory response compared to antibiotic treatment in mice. Antimicrobial Agents and Chemotherapy, 63(8), 2019. doi: 10.1128/AAC.00379-19.

Wendell H Fleming and Raymond W Rishel. Deterministic and stochastic optimal control. 1976.

Cesar O Flores, Justin R Meyer, Sergi Valverde, Lauren Farr, and Joshua S Weitz. Statistical structure of host–phage interactions. Proceedings of the National Academy of Sciences, 108(28):E288–E297, 2011.

Francesca Forti, Dwayne R. Roach, Marco Cafora, Maria E. Pasini, David S. Horner, Ersilia V. Fiscarelli, Martina Rossitto, Lisa Cariani, Federica Briani, Laurent Debarbieux, and Daniela Ghisotti. Design of a broad-range bacteriophage cocktail that reduces pseudomonas aeruginosa biofilms and treats acute infections in two animal models. Antimicrobial Agents and Chemotherapy, 62(6), 2018. ISSN 0066-4804. doi: 10.1128/AAC.02573-17.

Hiroshi Fukuhara, Yasushi Ino, and Tomoki Todo. Oncolytic virus therapy: a new era of cancer treatment at dawn. Cancer science, 107(10):1373–1379, 2016.

Andrzej Gorski, Ryszard Miedzybrodzki, Jan Borysowski, Krystyna Dabrowska, Piotr Wierzbicki, Monika Ohams, Grazyna Korczak-Kowalska, Natasza Olszowska-Zaremba, Marzena Lusiak-Szelachowska, Marlena Kłak, et al. Phage as a modulator of immune responses: practical implications for phage therapy. In Advances in virus research, volume 83, pages 41–71. Elsevier, 2012.

MT Hale, Y Wardi, H Jaleel, and M Egerstedt. Hamiltonian-based algorithm for optimal control. arXiv preprint arXiv:1603.02747, 2016.

Negar Hashemian and Antonios Armaou. Stochastic mpc design for a two-component granulation process. In 2017 American Control Conference (ACC), pages 4386–4391. IEEE, 2017.

Katarzyna Hodyra-Stefaniak, Paulina Miernikiewicz, Jarosław Drapała, Marek Drab, Ewa Jończyk-Matysiak, Dorota Lecion, Zuzanna Kaźmierczak, Weronika Beta, Joanna Majewska, Marek Harhala, et al. Mammalian host-versus-phage immune response determines phage fate in vivo. Scientific reports, 5:14802, 2015.

Taesoo Jang, Hee-Dae Kwon, and Jeehyun Lee. Free terminal time optimal control problem of an hiv model based on a conjugate gradient method. Bulletin of mathematical biology, 73(10):2408–2429, 2011.

Patrick Jault, Thomas Leclerc, Serge Jennes, Jean Paul Pirnay, Yok-Ai Que, Gregory Resch, Anne Françoise Rousseau, François Ravat, Hervé Carsin, Ronan Le Floch, Jean Vivien Schaal, Charles Soler, Cindy Fevre, Isabelle Arnaud, Laurent Bretaudeau, and Jérôme Gabard. Efficacy and tolerability of a cocktail of bacteriophages to treat burn wounds infected by pseudomonas aeruginosa (phagoburn): a randomised, controlled, double-blind phase 1/2 trial. The Lancet Infectious Diseases, 19(1):35 – 45, 2019. ISSN 1473-3099.

Serge Jennes, Maia Merabishvili, Patrick Soentjens, Kim Win Pang, Thomas Rose, Elkana Keersebilck, Olivier Soete, Pierre-Michel Françcois, Simona Teodorescu, Gunther Verween, et al. Use of bacteriophages in the treatment of colistin-only-sensitive pseudomonas aeruginosa septicaemia in a patient with acute kidney injury—a case report. Critical Care, 21(1):129, 2017.

Carl H June, Roddy S O’Connor, Omkar U Kawalekar, Saba Ghassemi, and Michael C Milone. Car t cell immunotherapy for human cancer. Science, 359(6382):1361–1365, 2018.

Kaitlyn E. Kortright, Benjamin K. Chan, Jonathan L. Koff, and Paul E. Turner. Phage therapy: A renewed approach to combat antibiotic-resistant bacteria. Cell Host & Microbe, 25(2):219 – 232, 2019. ISSN 1931-3128. doi: https://doi.org/10.1016/j.chom.2019.01.014.

Elizabeth Martin Kutter, Sarah J Kuhl, and Stephen T Abedon. Re-establishing a place for phage therapy in western medicine. Future microbiology, 10(5):685–688, 2015.

Sean E Lawler, Maria-Carmela Speranza, Choi-Fong Cho, and E Antonio Chiocca. Oncolytic viruses in cancer treatment: a review. JAMA oncology, 3(6):841–849, 2017.

Urszula Ledzewicz, Mohammad Naghnaeian, and Heinz Schäattler. Optimal response to chemotherapy for a mathematical model of tumor–immune dynamics. Journal of mathematical biology, 64(3):557–577, 2012.

Chung Yin (Joey) Leung and Joshua S. Weitz. Modeling the synergistic elimination of bacteria by phage and the innate immune system. Journal of Theoretical Biology, 429:241 – 252, 2017. ISSN 0022-5193.

William Manzanares, Margot Lemieux, Pascal L Langlois, and Paul E Wischmeyer. Probiotic and synbiotic therapy in critical illness: a systematic review and meta-analysis. Critical care, 20(1):262, 2016.

Shawna McCallin, Jessica C. Sacher, Jan Zheng, and Benjamin K. Chan. Current state of compassionate phage therapy. Viruses, 11(4), 2019. ISSN 1999-4915. doi: 10.3390/v11040343.

Carl R Merril, Dean Scholl, and Sankar L Adhya. The prospect for bacteriophage therapy in western medicine. Nature Reviews Drug Discovery, 2(6):489, 2003.

Rachael L Miller Neilan, Elsa Schaefer, Holly Gaff, K Renee Fister, and Suzanne Lenhart. Modeling optimal intervention strategies for cholera. Bulletin of mathematical biology, 72(8):2004–2018, 2010.

JIM O’neill. Antimicrobial resistance: tackling a crisis for the health and wealth of nations. Review on antimicrobial resistance, 1(1):1–16, 2014.

Elizabeth A Parker, Tina Roy, Christopher R D’Adamo, and L Susan Wieland. Probiotics and gastrointestinal conditions: An overview of evidence from the cochrane collaboration. Nutrition, 45:125–134, 2018.

Rafael Peña-Miller, David Lähnemann, Hinrich Schulenburg, Martin Ackermann, and Robert Beardmore. Selecting against antibiotic-resistant pathogens: optimal treatments in the presence of commensal bacteria. Bulletin of mathematical biology, 74(4):908–934, 2012.

Lev Semenovich Pontryagin, EF Mishchenko, VG Boltyanskii, and RV Gamkrelidze. The mathematical theory of optimal processes. 1962.

Dwayne R Roach, Chung Yin Leung, Marine Henry, Eric Morello, Devika Singh, James P Di Santo, Joshua S Weitz, and Laurent Debarbieux. Synergy between the host immune system and bacteriophage is essential for successful phage therapy against an acute respiratory pathogen. Cell host & microbe, 22(1):38–47, 2017.

Robert Rowthorn and Selma Walther. The optimal treatment of an infectious disease with two strains. Journal of mathematical biology, 74(7):1753–1791, 2017.

Shafiqul Alam Sarker, Shamima Sultana, Gloria Reuteler, Deborah Moine, Patrick Descombes, Florence Charton, Gilles Bourdin, Shawna McCallin, Catherine Ngom-Bru, Tara Neville, et al. Oral phage therapy of acute bacterial diarrhea with two coliphage preparations: a randomized trial in children from bangladesh. EBioMedicine, 4:124–137, 2016.

Robert T Schooley, Biswajit Biswas, Jason J Gill, Adriana Hernandez-Morales, Jacob Lancaster, Lauren Lessor, Jeremy J Barr, Sharon L Reed, Forest Rohwer, Sean Benler, et al. Development and use of personalized bacteriophage-based therapeutic cocktails to treat a patient with a disseminated resistant acinetobacter baumannii infection. Antimicrobial agents and chemotherapy, 61(10):e00954–17, 2017.

Josef Stoer and Roland Bulirsch. Introduction to numerical analysis, volume 12. Springer Science & Business Media, 2013.

Y Tanji, T Shimada, M Yoichi, K Miyanaga, K Hori, and H Unno. Toward rational control of escherichia coli o157: H7 by a phage cocktail. Applied Microbiology and Biotechnology, 64(2):270–274, 2004.

Jeremy J Thibodeaux and Timothy P Schlittenhardt. Optimal treatment strategies for malaria infection. Bulletin of mathematical biology, 73(11):2791–2808, 2011.

Yorai Wardi, Magnus Egerstedt, and Muhammad Umer Qureshi. Hamiltonian-based algorithm for relaxed optimal control. In 2016 IEEE 55th Conference on Decision and Control (CDC), pages 7222–7227. IEEE, 2016.

Ry Young and Jason J Gill. Phage therapy redux-—what is to be done? Science, 350(6265):1163–1164, 2015.

Jiayi Zhang, Brittany L Kraft, Yanying Pan, Samantha K Wall, Anthea C Saez, and Paul D Ebner. Development of an anti-salmonella phage cocktail with increased host range. Foodborne pathogens and disease, 7(11):1415–1419, 2010.

